# Herbivore feeding behavior validates optimal defense theory for specialized metabolites within plants

**DOI:** 10.1101/2021.06.27.450062

**Authors:** Pascal Hunziker, Sophie Konstanze Lambertz, Konrad Weber, Christoph Crocoll, Barbara Ann Halkier, Alexander Schulz

## Abstract

Numerous plants protect themselves from attackers using specialized metabolites. The biosynthesis of these deterrent, often toxic metabolites is costly, as their synthesis diverts energy and resources on account of growth and development. How plants diversify investments into growth and defense is explained by the optimal defense theory. The central prediction of the optimal defense theory is that plants maximize growth and defense by concentrating specialized metabolites in tissues that are decisive for fitness. To date, supporting physiological evidence merely relies on the correlation between plant metabolite distribution and animal feeding preference. Here, we use glucosinolates as a model to examine the effect of changes in chemical defense distribution on actual feeding behavior. Taking advantage of the uniform glucosinolate distribution in transporter mutants, we show that high glucosinolate accumulation in tissues important to fitness protects them by guiding larvae of a generalist herbivore to feed on other tissues. Moreover, we show that mature leaves of *Arabidopsis thaliana* supply young leaves with glucosinolates to optimize defense against herbivores. Our study provides physiological evidence for the central hypothesis of the optimal defense theory and sheds light on the importance of integrating glucosinolate biosynthesis and transport for optimizing plant defense.

---

Spatial and temporal variation of metabolically costly defenses within a plant is rationalized by the optimal defense theory, which predicts that defenses are allocated based on the fitness value and vulnerability of individual plant parts to simultaneously optimize growth and defense ^1–3^. Most research into plant defense theory is focused on its ecological-evolutionary aspects, which addresses one aspect of the optimal defense theory ^4,5^. Physiological studies addressing optimal defense within individual plants exclusively rely on altered animal feeding behavior on plants that are e.g. genetically engineered to lack biosynthesis or activation of defensive metabolites ^6,7^. Due to the lack of genetic tools to manipulate chemical defense distribution, empirical studies addressing the effect of optimal defense distribution within a plant on animal feeding preference and behavior are still missing. Here, we asked how the allocation of chemical defense compounds affects herbivore feeding behavior using glucosinolates in *Arabidopsis thaliana* as a model.

Glucosinolates are amino acid-derived thio-glucosides that yield herbivore-deterrent catabolites upon tissue damage-triggered enzymatic hydrolysis by myrosinases ^8^. However, many herbivores circumvent glucosinolate-based defenses via diverse behavioral and physiological counter-adaptations ^9^. High selective pressure by coevolving predators conserved the production of glucosinolates despite large energy costs ^10,11^. *A. thaliana* meets the predictions of the optimal defense theory: young leaves are more valuable than old ones and display much higher glucosinolate concentration ^12,13^. Similar ontogenetic patterns were reported for many other species of the Brassicales order, indicating existence of a universal glucosinolate distribution pattern ^14–18^. To examine the glucosinolate distribution in rosettes of the reference plant *A. thaliana*, we grouped leaves of 5-week-old plants into three cohorts according to their age (**Fig. 1a**). Liquid chromatography-mass spectrometry revealed that the total glucosinolate concentration is highest in young leaves and decreases gradually with leaf age (**Fig. 1b**). To investigate whether the ontogenetic pattern of glucosinolate concentrations across the rosette correlate with transcript levels of key enzymes involved in glucosinolate biosynthesis, we measured the expression of *CYP79B2, CYP83A1, MAM1, MAM3, SUR1* and *UGT74C1* by quantitative reverse transcription polymerase chain reaction. Gene expression was lowest in young leaves and gradually increased with leaf age, suggesting that glucosinolates are primarily produced in old leaves and subsequently transported to young ones (**Fig. 1c**). To test this hypothesis, we analyzed the tissue-specific glucosinolate distribution in knockout mutants of genes encoding the well-characterized H^+^/glucosinolate symporters NPF2.10/GTR1, NPF2.11/GTR2 and NPF2.9/GTR3 ^19,20^. In *gtr1 gtr2* double knockout mutants, the gradient of total glucosinolates across leaf groups was abolished, demonstrating that GTR1/GTR2-mediated transport establishes the ontogenetic glucosinolate distribution in the rosette (**Fig. 1b**). Compared to wild-type, total glucosinolate levels were significantly reduced in young leaves and increased in old and mature ones, indicating that GTR1 and GTR2 are responsible for the export of glucosinolates from old leaves. Intriguingly, indolic glucosinolates – particularly 1-methoxy-indol-3-ylmethyl glucosinolate – retained a gradual distribution in *gtr1 gtr2* (**Supplemental Fig. 1**). Analysis of the tissue-specific glucosinolate distribution in *gtr1 gtr2 gtr3* triple knockout mutants – GTR3 has high affinity towards indolic glucosinolates – showed that high indolic glucosinolate levels in young leaves are still maintained in absence of all three glucosinolate transporters, despite significant accumulation in old and mature leaves leading to their uniform distribution among leaf cohorts (**Supplemental Fig. 1**). To test whether tissue-specific induction of biosynthesis compensates for the absence of glucosinolate supply from old and mature leaves, we analyzed the spatial gene expression of key biosynthetic enzymes in *gtr1 gtr2* and found significant genotype × tissue interactions for *CYP79B2, CYP83A1, MAM3* and *SUR1* (**Fig. 1c**). The transcript levels of these genes were reduced in old leaves, indicating that *GTR1/GTR2*-dependent over-accumulation of glucosinolates in old leaves may lead to feedback inhibition of glucosinolate biosynthesis gene expression (**Fig. 1b, c**). By contrast, expression of *CYP79B2* – a marker for indolic glucosinolate synthesis – was increased in young leaves suggesting that enzymes involved in indolic glucosinolate biosynthesis are *GTR1/GTR2*-dependently repressed in young leaves. This supports that the altered glucosinolate distribution in *gtr1 gtr2* is caused by perturbation of transport that cannot be entirely compensated for by biosynthesis.

**Fig. 1.**
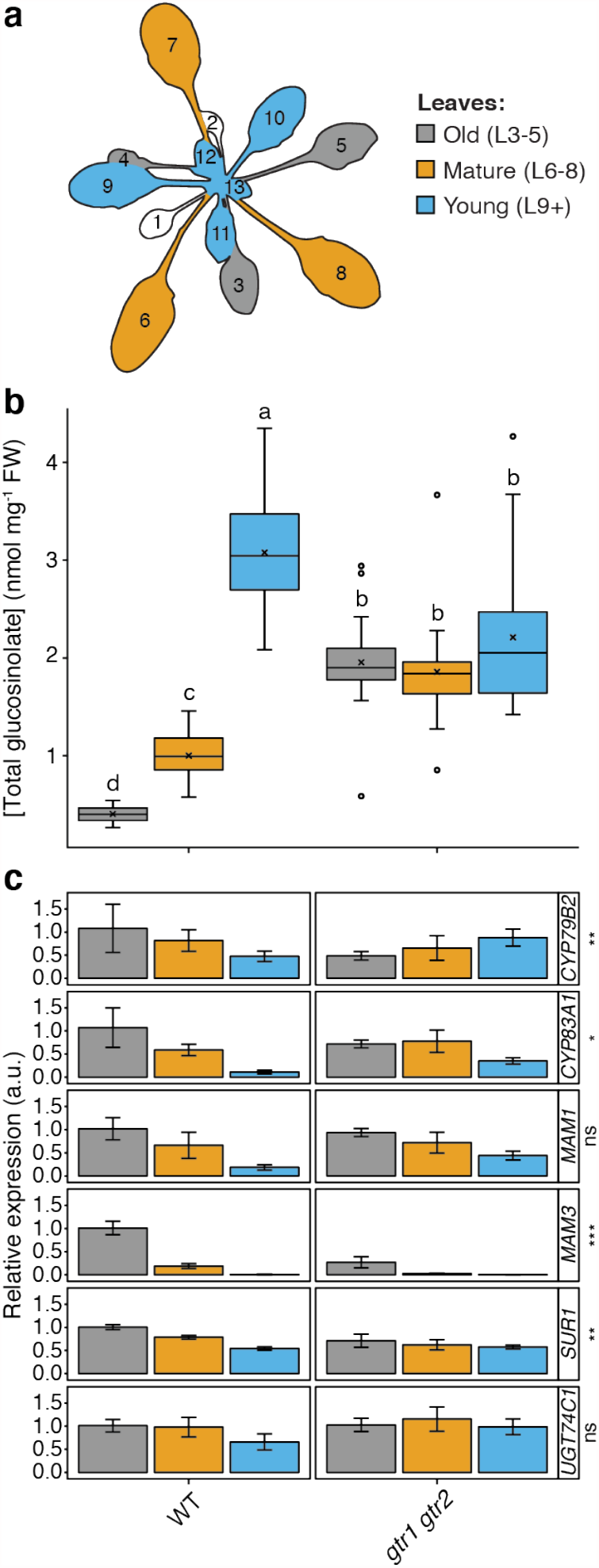
Tissue-specific analysis of glucosinolate concentrations and biosynthetic gene expression in transporter knockout mutants. **a**, Harvesting scheme depicting the silhouette of an *Arabidopsis thaliana* rosette used for sampling. Leaves are numbered according to ontogenetic development starting with the first true leaf. Colors indicate the grouping into cohorts of old (L3-5; grey), mature (L6-8; orange) and young leaves (L9+; blue). **b**, Tissue-specific total glucosinolate concentration of wild-type (WT) and *gtr1 gtr2* double knockout transporter mutants. Colors indicate leaf cohorts according to (A). Letters indicate significant differences of genotype × tissue interactions (Two-way ANOVA with post-hoc Tukey HSD test; n = 10; p<0.001). Pooled data from two independent replicates are shown. **c**, Expression analysis of glucosinolate biosynthetic genes in WT and *gtr1gtr2* mutants. Colors indicate leaf cohorts according to (A). Transcript levels of *CYP79B2, CYP83A1, MAM1, MAM3, SUR1* and *UGT74C1* are shown relative to the abundance in old leaves (L3-5) of the wild-type. Values represent means ± SD. Stars indicate significance of genotype × tissue interactions for each gene (Two-way ANOVA; n = 4; ns, not significant; *p<0.05; **p<0.01; ***p<0.001).

Next, we leveraged the uniform distribution of glucosinolates in the rosette of *gtr1 gtr2* mutants and used glucosinolate-based chemical defense as a model to explore the effect of defense distribution on herbivore feeding behavior. We performed a bioassay using larvae of *Spodoptera littoralis*, a generalist insect herbivore that infests plants from at least 40 families including the families of the Brassicales order and many economically important species ^21^. *S. littoralis* showed a strong preference (100 %, n = 90) for feeding on old and mature leaves of the wild-type, as *S. littoralis* larvae caused severe feeding damage on old and mature leaves while avoiding the young ones (**Fig. 2a, b, Supplemental Fig. 2**). This feeding behavior inversely correlates with leaf glucosinolate concentrations and is consistent with the predictions of the optimal defense theory for defense distribution within the plant. Strikingly, we found that the feeding preference is inversed (98.5 %, n = 67) on *gtr1 gtr2* (**Fig. 2a, b**). Thus, the spatial pattern of glucosinolates distribution governs the feeding behavior of *S. littoralis*. This conclusion is further supported by the observation of a preference for young leaves (100 %, n = 56) on glucosinolate-deficient *myb28 myb29 cyp79b2 cyp79b3* quadruple mutants (**Fig. 2b, Supplemental Fig. 3**). As *GTR1* and *GTR2* expression is not affected in the quadruple mutant ^22^, the finding demonstrates that the feeding behavior specifically depends on glucosinolates, thus eliminating a potential role of GTR1/GTR2-mediated distribution of other metabolites (e.g. nutrients or hormones). Over-accumulation of glucosinolates in old leaves of *gtr1 gtr2* cannot explain the feeding behavior, as larvae also avoided feeding on old leaves of glucosinolate-deficient plants, probably due to the relative nutritional value and digestibility (**Figs. 1b, 2b**). The feeding behavior on biosynthetic and transport-deficient mutants supports that high glucosinolate concentrations in young leaves are crucial to their protection. Consistent with previous studies, showing that glucosinolates delay larval development of *S. littoralis* due to energy-consumptive detoxification ^23^, larvae gained significantly more weight on glucosinolate-deficient plants compared to wild-type (**Fig. 2c**). By contrast, *S. littoralis* feeding on glucosinolate transporter mutants did not affect the weight of larvae (**Fig. 2c**), possibly because the ingestion of the residual glucosinolates in young leaves of *gtr1 gtr2* is sufficient to delay growth compared to the glucosinolate-free young leaves of the quadruple mutant. Regardless, *A. thaliana* profits from transporter-mediated, non-uniform glucosinolate distribution in the rosette, which prevents herbivore attacks against the most valuable and vulnerable tissues, thus providing physiological evidence for the central hypothesis of the optimal defense theory.

**Fig. 2.**
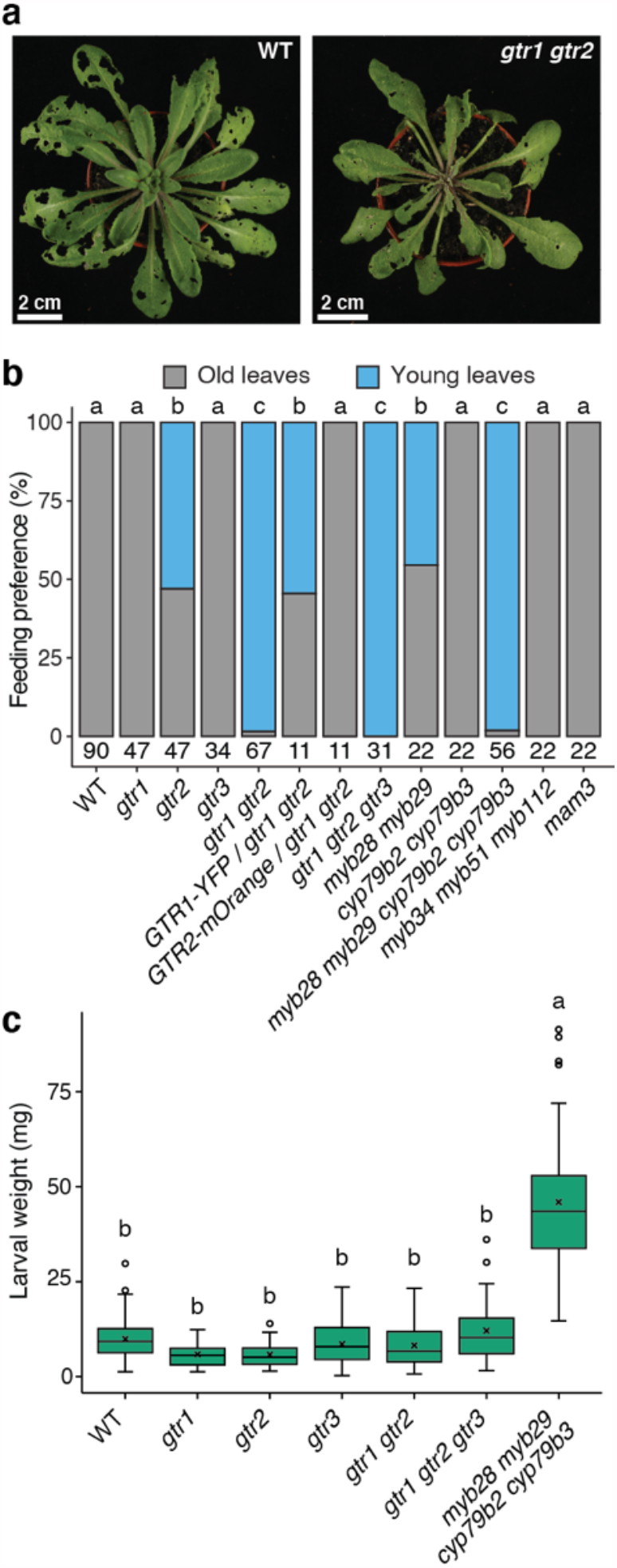
*GTR1* and *GTR2* determine the feeding preference of larvae of generalist herbivore *Spodoptera littoralis*. **a**, Photographs of 5-week-old wild-type (WT) and *gtr1gtr2* double knockout mutant plants imaged 8 days after infection with newly hatched larvae in a non-choice experimental set-up. Note that the young leaves in the center of the rosette are almost completely consumed in *gtr1 gtr2*. Scale bar = 2 cm. **b**, Quantification of *S. littoralis* feeding pattern on wild-type (WT), glucosinolate transporter mutants (*gtr1, gtr2, gtr3, gtr1 gtr2, gtr1 gtr2 gtr3, GTR1-YFP/gtr1 gtr2* and *GTR2-mOrange/gtr1 gtr2*) and glucosinolate biosynthesis mutants (*myb28 myb29, cyp79b2 cyp79b3, myb28 myb29 cyp79b2 cyp79b3, myb34 myb51 myb112* and *mam3*). Number of biological replicates (n) is indicated below stacked bars. Pooled data from one to eight independent replicates are shown. Letters above stacked bars indicate significant differences between genotypes (Fisher’s exact test, p<0.05). **c**, Quantification of larval weight of *S. littoralis* following 8 days of feeding on 4-week-old wild-type and mutant plants. Letters depict significant differences between genotypes (Two-way ANOVA with post-hoc Tukey HSD test; n = 103; 30; 33; 38; 83; 36; 44; p<0.001). Pooled data from three independent replicates are shown.

In addition to defense allocation within the plant, the optimal defense theory also predicts that defense is increased upon attack. To test how the tissue-specific distribution of glucosinolates is affected by the plant-herbivore interaction, we measured their concentration in young wild-type leaves upon *S. littoralis* feeding. As young leaves of the mutants were completely consumed during bioassays, we further used mechanical wounding to mimic generalist herbivores. Indolic glucosinolate levels were significantly (∼3-fold) induced by both treatments, indicating that herbivore-associated molecular patterns are negligible for this response (**Fig. 3**). These increases also occurred in *gtr1 gtr2* and *gtr1 gtr2 gtr3*, suggesting tissue-specific induction of *de novo* indolic glucosinolate biosynthesis upon wounding (**Fig. 3b, Supplemental Fig. 5**), which is consistent with the reported upregulation of genes involved in indolic glucosinolate biosynthesis upon *S. littoralis* feeding and wounding ^24^. Furthermore, *S. littoralis* exhibited a feeding preference for young leaves on aliphatic glucosinolate-deficient *myb28 myb29*, and not indolic glucosinolate-deficient *cyp79b2 cyp79b3* and *myb34 myb51 myb122* mutants, demonstrating that aliphatic and not indolic glucosinolates are critical for establishing the feeding preference (**Fig. 2b, Supplemental Fig. 3**). Indeed, the levels of long-chain aliphatic glucosinolates in young leaves increased significantly upon both feeding and wounding (**Supplemental Figs. 4, 5**).

**Fig. 3.**
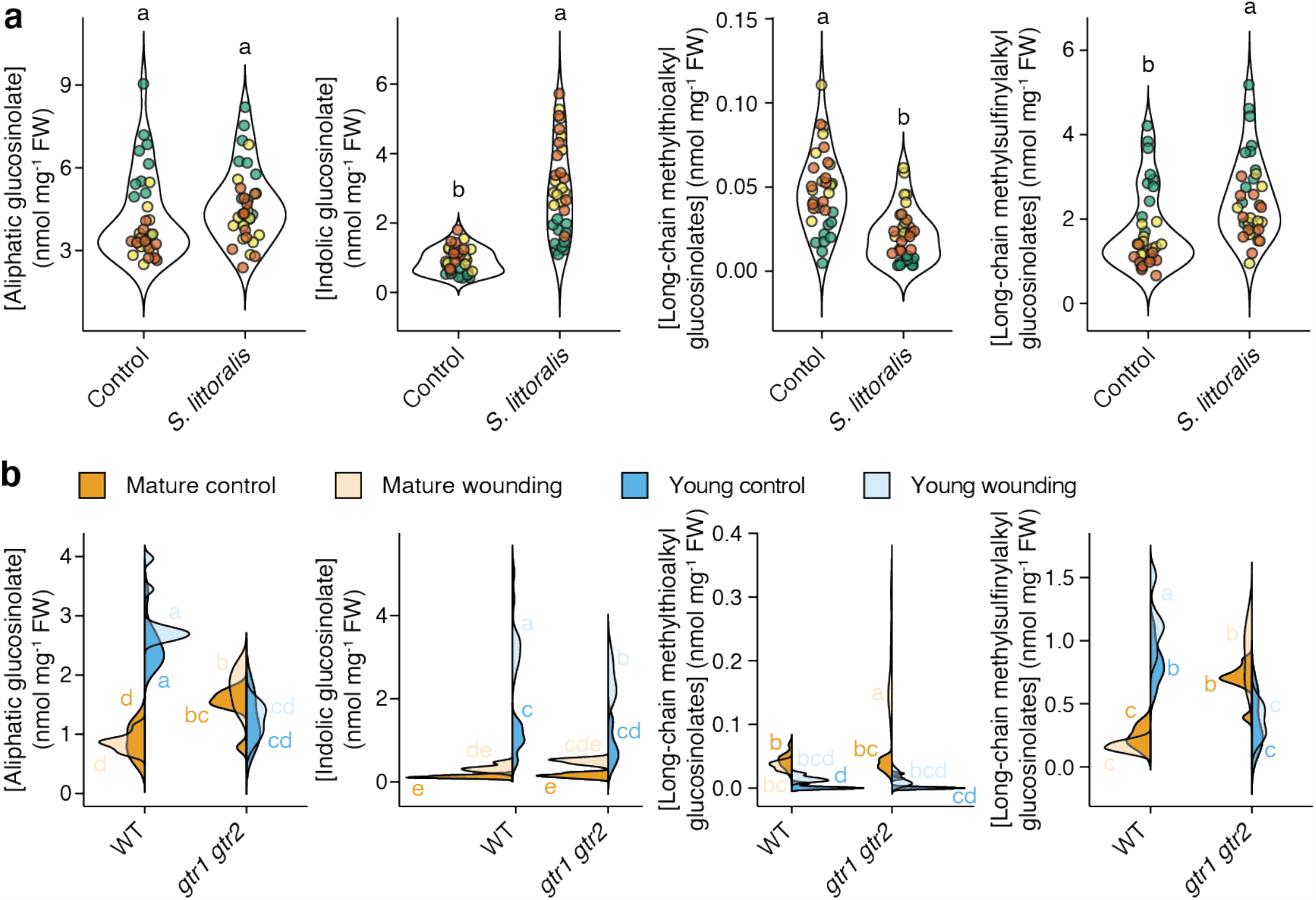
Tissue-specific accumulation of glucosinolates upon mechanical wounding and herbivore attack. **a**, Perturbations of aliphatic, indolic, long-chain methylthioalkyl and long-chain methylsulfinylalkyl glucosinolate concentrations in young leaves (L9+) of wild-type plants following eight days of *Spodoptera littoralis* feeding. Letters depict significant differences among treatments (Two-way ANOVA with post-hoc Tukey HSD test; n = 11; p<0.001). Pooled data from three independent replicates are shown. Data points represent biological replicates and are color-coded according to experimental replicates. **b**, Tissue-specific perturbations of aliphatic, indolic, long-chain methylthioalkyl and long-chain methylsulfinylalkyl glucosinolate concentrations in mature (L6-8; orange) and young (L9+; blue) leaves of wild-type and *gtr1 gtr2* double knockout mutants upon repetitive mechanical wounding of old leaves (L3-5). Letters indicate significant differences of genotype × tissue interactions (Two-way ANOVA with post-hoc Tukey HSD test, n = 10, p<0.001). Leaves were grouped according to Fig. 1a.

Moreover, specifically long-chain methylsulfinylalkyl glucosinolates accumulate, indicating that feeding-inflicted wounding triggers *S*-oxygenation on the side-chain in young leaves (**Fig. 3, Supplemental Figs. 4, 5**). *S*-oxygenation in response to wounding was attenuated in *gtr1 gtr2* and, interestingly, the mutant over-accumulated the corresponding precursors (i.e. methylthioalkyl glucosinolates) in unwounded mature leaves (**Fig. 3b, Supplemental Fig. 5**). Yet, the accumulation and modification of long-chain aliphatic glucosinolates is not required for the defense of young leaves, as *S. littoralis* showed a habitual feeding preference for old leaves on long-chain aliphatic glucosinolate-deficient *mam3* mutants (**Fig. 2b, Supplemental Fig. 3**). Hence, basal levels of total aliphatic glucosinolates are critical for optimal defense of young tissues against generalist herbivores such as *S. littoralis*.

Based on our results, we propose a model that predicts a new function for mature tissues in serving as relays that orchestrate glucosinolate biosynthesis and transport to protect tissues with higher fitness values from attack (**Supplemental Fig. 6**). Briefly, glucosinolates are translocated from root to shoot via the xylem, which is controlled by a GTR1/GTR2/GTR3-mediated retention mechanism that is inactivated upon feeding-induced wounding. This is supported by our finding that mature leaves of untreated *gtr1 gtr2 gtr3* mutants display elevated long-chain methylthioalkyl glucosinolate concentrations, which could not be further induced by wounding (**Supplemental Fig. 5**). In mature leaves, glucosinolates are GTR1/GTR2-dependently loaded into the phloem for transport to young leaves, which is supported by the expression of *GTR1* and *GTR2* in the vasculature and hydathodes (**Supplemental Fig. 7**). Young, immature plant tissues are well-known to be strong sinks for primary metabolites that are supplied by mature tissues. Due to metabolic constraints, biosynthesis of specialized metabolites – such as glucosinolates – in mature leaves may thus be more adaptive for plants than producing them in young leaves, which is consistent with the sub-hypothesis of the optimal defense theory predicting a trade-off between defense and other plant functions such as growth and reproduction. Hence, using glucosinolate-based defense of *A. thaliana* against the generalist herbivore *S. littoralis*, we were able to provide experimental evidence for the physiological aspects of the optimal defense theory and highlight the importance of coordinating biosynthesis and transport processes of defensive metabolites to obtain most favorable distribution within a plant for optimizing growth and defense.

## Acknowledgments

We thank D. G. Vassão and V. Jeschke for providing *S. littoralis* eggs. This work was funded by the Danish National Research Foundation (grant DNRF99).

## Author contributions

P.H. and S.K.L. designed the study. P.H., S.K.L., K.W. and C.C. performed experiments and analyzed data. B.A.H. and A.S. provided funding, resources and supervision. P.H. drafted the manuscript. All authors discussed the results and contributed to the manuscript.

## Methods

### Biological material

*Arabidopsis thaliana* (L.) Heynh. plants were grown on soil in a walk-in climate chamber under short-day conditions (10 h light, 150 µmol m^-2^ s^-1^, 21 °C, 70 % humidity). The Columbia-0 accession (Col-0) was used as wild-type. The following, previously described T-DNA insertion and transgenic lines were used: *myb28 myb29* ^25^, *cyp79b2 cyp79b3* ^26^, *myb28 myb29 cyp79b2 cyp79b3* ^22^, *myb34 myb51 myb122* ^27^, *mam3* ^28^, *gtr1* ^20^, *gtr2* ^20^, *gtr3* ^19^, *gtr1 gtr2* ^20^, *gtr1 gtr2 gtr3* ^19^, *pGTR1::NLS-GUS-GFP* ^20^, *pGTR2::GTR2-NLS-GUS-GFP* ^20^, *pGTR1::GTR1-YFP* in *gtr1 gtr2* ^20^ and *pGTR2::GTR2-mOrange2* in *gtr1 gtr2* ^20^. Egg clutches of *Spodoptera littoralis* Boisduval (African cotton leafworm) were a generous gift from Daniel G. Vassão of the Max Planck Institute for Chemical Ecology (Jena, Germany), and reared on an artificial diet based on white beans at 18-20°C under natural light ^23,29^.

### Desulfo-glucosinolate analysis by LC-MS

Leaves of 4-week-old plants were harvested according to the harvesting scheme presented in **Fig. 1a** and immediately submerged in 85 % methanol containing 50 µM *p*-hydroxybenzyl glucosinolate as internal standard. Desulfo-glucosinolates were purified as described previously ^30^. Samples were 10-fold diluted with deionized water prior to analysis by liquid chromatography coupled to mass spectrometry. Chromatography was performed on an Advance UHPLC system (Bruker, Bremen, Germany). Separation was achieved on a Kinetex 1.7u XB-C18 column (100 × 2.1 mm, 1.7 µm, 100 Å, Phenomenex, Torrance, CA, USA). Formic acid (0.05 %) in water and acetonitrile (supplied with 0.05 % formic acid) were employed as mobile phases A and B respectively. The elution profile was: 0-0.5 min, 2 % B; 0.5-1.2 min, 2-30 % B; 1.2-2.0 min 30-100 % B, 2.0-2.5 min 100 % B, 2.5-2.6 min, 100-2 % B and 2.6-4 min 2 % B. The mobile phase flow rate was 400 µl min^-1^. The column temperature was maintained at 40°C. The liquid chromatography was coupled to an EVOQ Elite TripleQuad mass spectrometer (Bruker, Bremen, Germany) equipped with an electrospray ion source (ESI) operated in positive ionization mode. The instrument parameters were optimized by infusion experiments with pure standards. The ion spray voltage was maintained at +3500 V. Cone temperature was set to 300°C and cone gas to 20. Heated probe temperature was set to 400 °C and probe gas flow to 40 psi. Nebulizing gas was set to 60 psi and collision gas to 1.5 mTorr. Nitrogen was used as probe and nebulizing gas and argon as collision gas. Active exhaust was constantly on. Multiple reaction monitoring (MRM) was used to monitor analyte precursor ion → product ion transitions. Detailed values for mass transitions can be found ^31,32^. Both Q1 and Q3 quadrupoles were maintained at unit resolution. Bruker MS Workstation software (Version 8.21, Bruker, Bremen, Germany) was used for data acquisition and processing. Linearity in ionization efficiencies were verified by analyzing dilution series. Glucosinolate concentrations were normalized to fresh weight. Glucosinolates were grouped as follows: Aliphatic (3-methylthiopropyl, 3-methylsulfinylpropyl, 4methylthiobutyl, 4-methylsulfinylbutyl, 5-methylsulfinylpentyl, 7-methylthioheptyl, 7-methylsulfinylheptyl, 8-methylthiooctyl and 8-methylsulfinyloctyl glucosinolate), indolic (indol-3-ylmethyl, 1-methoxy-indol-3-ylmethyl and 4-methoxy-indol-3-ylmethyl glucosinolate), short-chain aliphatic (3-methylthiopropyl, 3-methylsulfinylpropyl, 4methylthiobutyl, 4-methylsulfinylbutyl, 5-methylsulfinylpentyl glucosinolate), long-chain aliphatic (7-methylthioheptyl, 7-methylsulfinylheptyl, 8-methylthiooctyl and 8-methylsulfinyloctyl glucosinolate), methylthioalkyl (3-methylthiopropyl, 4methylthiobutyl, 7-methylthioheptyl and 8-methylthiooctyl glucosinolate) and methylsulfinylalkyl (3-methylsulfinylpropyl, 4-methylsulfinylbutyl, 5-methylsulfinylpentyl, 7-methylsulfinylheptyl and 8-methylsulfinyloctyl glucosinolate) glucosinolates.

### GUS staining

Plant material was collected in 90% acetone, incubated for at least 20 min and washed with 50 mM sodium phosphate buffer pH 7.0 for 5-10 min. The rinsing solution was replaced with staining mix (50 mM sodium phosphate buffer pH 7.0, 0.1% Triton X-100, 0.5 mM K4Fe(CN)6, 0.5 mM K3Fe(CN)6, 0.5 mg/ml X-Gluc). Plant material was then vacuum-infiltrated for 5 min at room temperature and incubated at 37°C in the dark overnight. The reaction was stopped by replacing the staining solution with 70% ethanol, which was exchanged several times until the tissue was completely cleared. Images were taken using a Leica M205 FA stereomicroscope (Leica Microsystems, Heerbrugg, Switzerland) or an Epson Perfection V550 Photo color scanner (Seiko Epson Corporation, Suwa, Nagano, Japan). White balance was adjusted using Photoshop CC (Version 19.0, Adobe Systems Inc., San Jose, CA, USA).

### Insect bioassays

Neonate *Spodoptera littoralis* were placed on 3,5- to 4-week-old plants. Plant genotypes were contained in separate plastic boxes. In each box, eleven plants were distributed and three larvae were placed at the center of the rosette of each individual plant. Larvae were able to move between plants within each box and allowed to feed for 7-8 days. The mass of individual larvae was determined by a Mettler Toledo PG503-S DeltaRange® precision balance (Mettler Toledo, Columbus, Ohio, USA).

### RNA extraction, cDNA synthesis and RT-qPCR

For total RNA extraction, designated plant material of three plants was combined to one biological replicate and snap frozen in liquid nitrogen followed by homogenization to fine powder in a RETSCH bead-mill. RNA was extracted using Sigma Spectrum Plant Total RNA kit. Electrophoresis was used to assess native RNA quality on a 1% agarose gel. Next, 1 µg total RNA was DNase1-treated (Sigma) and used for cDNA synthesis (iScript™ from BIO RAD) with a blend of random hexamer and oligo(dT) primers in a 20 µl reaction. The obtained cDNA was 1:10 diluted in water and 2 µl were used in a 10 µl SYBR green (PowerUp SYBR™ applied biosystems) RT-qPCR reaction, run in a CFX384 Touch™ real-time PCR detection system from BIO RAD. RT-qPCR of genes of interest and the housekeeping transcript *UBC21* (At5g25760) were performed in standard cycling mode condition according to SYBR green manufacturer. Three technical replicates were run for each biological replicate. Relative expression was calculated with kinetic PCR efficiency correction and normalized to *UBC21* and further to WT sample “Old leaves (L3-5)”. Oligonucleotide sequences used in this study can be found in **Supplemental Table 1**.

### Mechanical wounding

Wounds were inflicted with forceps parallel to the longitudinal axis of the third, fourth and fifth leaves. The first wound was made at the leaf tip and the second wound was made so that it abutted the first, and so on until 40–50% of the leaf was wounded. The wounding was repeated three times in 3 h intervals. Unwounded plants served as a control and were randomly distributed among wounded plants. The plant material was harvested 24 h after the first round of wounding.

### Data visualization and statistics

Data visualization and statistical analyses were performed in R ^33^. Plots were produced using the ggplot2 package ^34^. Boxplots show median (center line), mean (cross), first quartile (lower hinge), third quartile (upper hinge), whiskers (extending 1.5 times the inter-quartile range) and possible outliers (circles). ANOVA analyses and post-hoc Tukey’s HSD test were performed using the agricolae and nmle packages ^35,36^. Prior to analysis, data were checked for normality and scedasticity using diagnostic plots. Glucosinolate concentrations were fitted to a linear model as a function of genotype × tissue × experimental replicate (**Fig. 1b, Supplemental Fig. 1**), treatment × experimental replicate (**Fig. 3a, Supplemental Fig. 4**) or genotype × treatment × tissue (**Fig. 3b, Supplemental Fig. 5**). Relative transcript levels were fitted to a linear model as a function of genotype × tissue (**Fig. 1c**). Larval weight was fitted to a linear model as a function of genotype × experimental replicate (**Fig. 2c, Supplemental Figs. 2b, 3b**). Additionally, glucosinolate concentrations were fitted to a mixed effect model accounting for box effects as a random factor (**Fig. 3a**). A likelihood ratio test was employed to compare linear and mixed effect models and revealed non-significant p-values for each independent variable. Based on this result, linear models were preferred over mixed effect models. Fisher’s exact test was performed using a Monte Carlo simulation with 10^7^ replicates. Post-hoc pairwise comparison with Bonferroni-adjusted p-values was performed using the rcompanion package ^37^. *F* values, p values and degrees of freedom can be found in **Supplemental Table 2, 3**.

## Supplemental information

**Supplemental Fig. 1**. Tissue-specific analysis of glucosinolates in wild-type (WT), *gtr1, gtr2, gtr3, gtr1 gtr2* and *gtr1 gtr2 gtr3* plants. Colors indicate the grouping of leaves according to the Fig. 1a. Letters indicate significant differences of genotype × tissue interactions (two-way ANOVA; n = 10; p<0.001). Pooled data from two independent replicates are shown for all genotypes except *gtr3*, which was solely included in one replicate.

**Supplemental Fig. 2**. Complementation of the *S. littoralis* feeding preference phenotype of *gtr1gtr2*. **a**, Photographs of 5-week-old wild-type (WT), *gtr1, gtr2, gtr1 gtr2, pGTR2:GTR2-mOrange/gtr1 gtr2* and *pGTR1:GTR1-YFP/gtr1 gtr2* mutant plants imaged 7 days after infection with newly hatched larvae in a non-choice experimental set-up. Scale bar = 2 cm. **b**, Quantification of larval weight of *S. littoralis* following 7 days of feeding on 4-week-old wild-type (WT) and mutant plants. Boxplots show median (center line), mean (cross), first quartile (lower hinge), third quartile (upper hinge), whiskers (extending 1.5 times the inter-quartile range) and possible outliers (circles). Letters depict significant differences between genotypes (two-way ANOVA with post-hoc Tukey HSD test; n = 36, 22, 40, 34, 29, 33; p<0.001).

**Supplemental Fig. 3**. *S. littoralis* feeding preference on loss-of-function mutants for glucosinolate biosynthesis. **a**, Photographs of 4-week-old wild-type (WT), *myb28 myb29, cyp79b2 cyp79b3, myb28 myb29 cyp79b2 cyp79b3, myb34 myb51 myb112* and *mam3* mutant plants imaged 7 days after infection with newly hatched larvae in a non-choice experimental set-up. Scale bar = 2 cm. **b**, Quantification of larval weight of *S. littoralis* following 7 days of feeding on 4-week-old wild-type and mutant plants. Boxplots show median (center line), mean (cross), first quartile (lower hinge), third quartile (upper hinge), whiskers (extending 1.5 times the inter-quartile range) and possible outliers (circles). Letters depict significant differences between genotypes (two-way ANOVA with post-hoc Tukey HSD test; n = 61; 61; 60; 59; 59; 30; p<0.001). Pooled data of two independent replicates are shown.

**Supplemental Fig. 4**. *S. littoralis*-induced perturbations of glucosinolate concentration in young leaves (L9+) of wild-type plants. Three newly hatched larvae per 5-week-old plant were allowed to feed for eight days. Young leaves were classified according to Fig. 1a. Asterisks indicate significant treatment effects (two-way ANOVA; n = 11; *p<0.05, **p<0.01, ***p<0.001). Pooled data from three independent experimental replicates are shown. Data points represent biological replicates and are color-coded according to experimental replicates.

**Supplemental Fig. 5**. Tissue-specific glucosinolate analysis in mature (L6-8; orange) and young (L9+; blue) leaves of wild-type (WT), *gtr1, gtr2, gtr1 gtr2* and *gtr1 gtr2 gtr3* upon repetitive mechanical wounding of old leaves (L3-5). Samples were harvested 24 hours after treatment. Letters indicate significant differences of genotype × tissue interactions (two-way ANOVA; n = 10; p<0.001). Leaves were grouped according to Fig. 1a.

**Supplemental Fig. 6**. Model for glucosinolate transport upon herbivore attack. **(1)** GTR1-3 prevent root-synthesized glucosinolates from translocation to the shoot via the xylem under normal growth conditions providing a retention mechanism ^19,38^. **(2)** Mechanical wounding of leaves by herbivore feeding attenuates GTR1/GTR2/GTR3-based glucosinolate retention in the root and leads to increased root-to-shoot translocation of mostly methylthioalkyl (3-methylthiopropyl, 4-methylthiobutyl, 7-methylthioheptyl and 8-methylthiooctyl) and indolic (indol-3-ylmethyl and 1-methoxy-indol-3-ylmethyl) glucosinolates via the xylem. Note that *GTR3* is exclusively expressed in roots ^39^. **(3)** GTR1 and GTR2 load root-derived and old leaf-synthesized glucosinolates into the phloem of old leaves. High accumulation of glucosinolates in the extracellular space inhibits glucosinolate biosynthesis in old leaves. **(4)** Glucosinolates are translocated towards carbon sinks via the phloem ^40–42^. **(5)** *S*-oxygenation of methylthioalkyl glucosinolates is induced either during transport in the phloem (i.e. inside the sieve element/companion cell-complex) or upon arrival in young leaves. Tissue-specific induction of *de novo* biosynthesis of indolic glucosinolates is independent of GTR1-3.

**Supplemental Fig. 7**. Gene expression patterns of *GTR1* and *GTR2* in 4-weeks-old plants. Scanning (**a-b**) and stereomicroscopic (**c-d**) images of GUS-stained transgenic plants that stably express *pGTR1::NLS-GUS-GFP* (**a**) and *pGTR2::NLS-GUS-GFP* (**b-d**). **c** and **d** represent higher magnification images of **b**. Scale bars = 1cm (a and b); 1 mm (c and d).

**Supplemental Table 1**. Oligonucleotides used for RT-qPCR analysis.

**Supplemental Table 2**. ANOVA tables for data shown in Figures.

**Supplemental Table 3**. ANOVA tables for data shown in Supplemental Figures.

