## Supplemental information for "Herbivore feeding behavior validates optimal defense theory for specialized metabolites within plants"

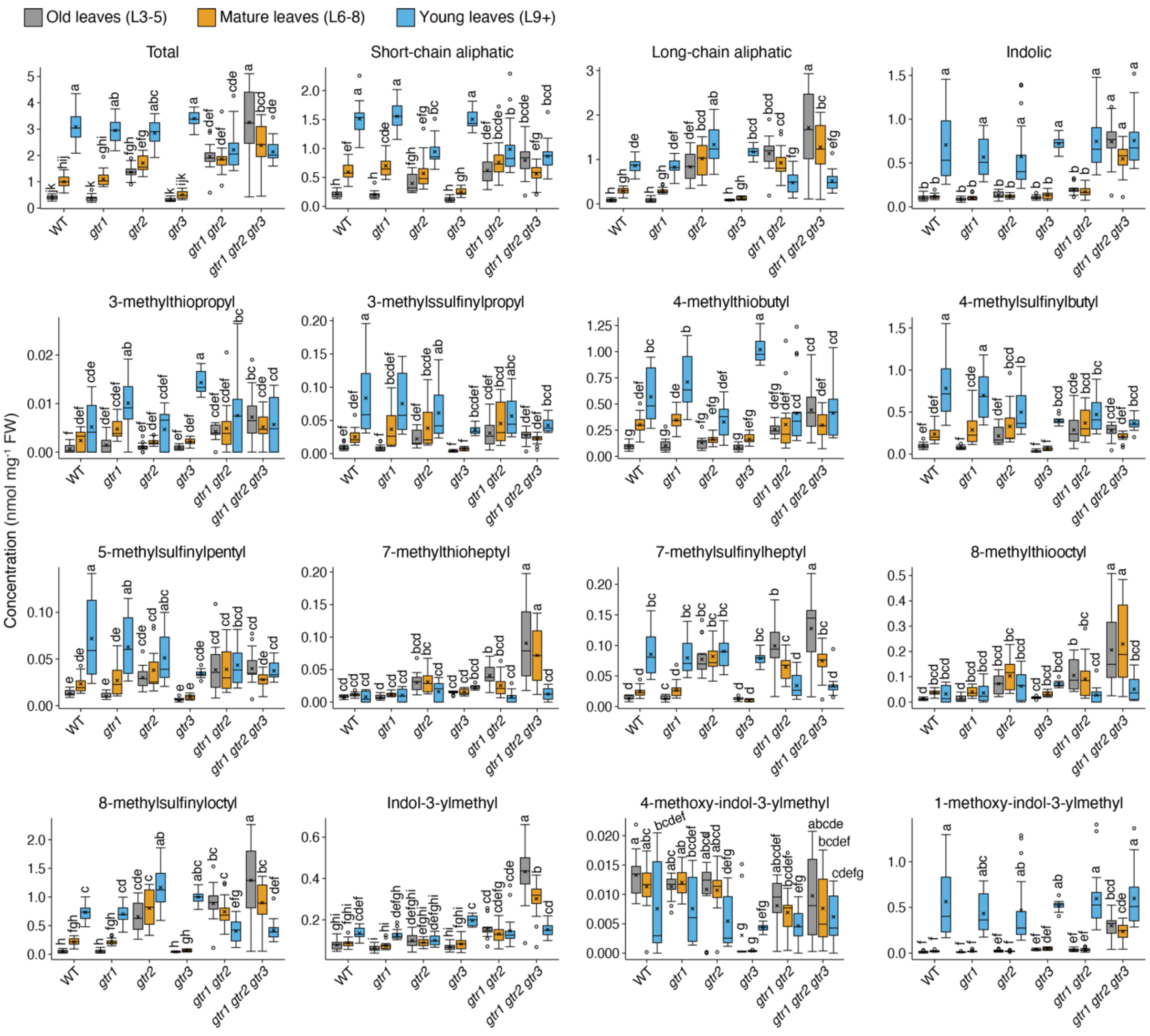


**Supplemental Fig. 1.** Tissue-specific analysis of glucosinolates in wild-type (WT), *gtr1*, *gtr2*, *gtr3*, *gtr1 gtr2* and *gtr1 gtr2 gtr3* plants. Colors indicate the grouping of leaves according to the Fig. 1a. Letters indicate significant differences of genotype × tissue interactions (two-way ANOVA; n = 10; p<0.001). Pooled data from two independent replicates are shown for all genotypes except *gtr3*, which was solely included in one replicate.

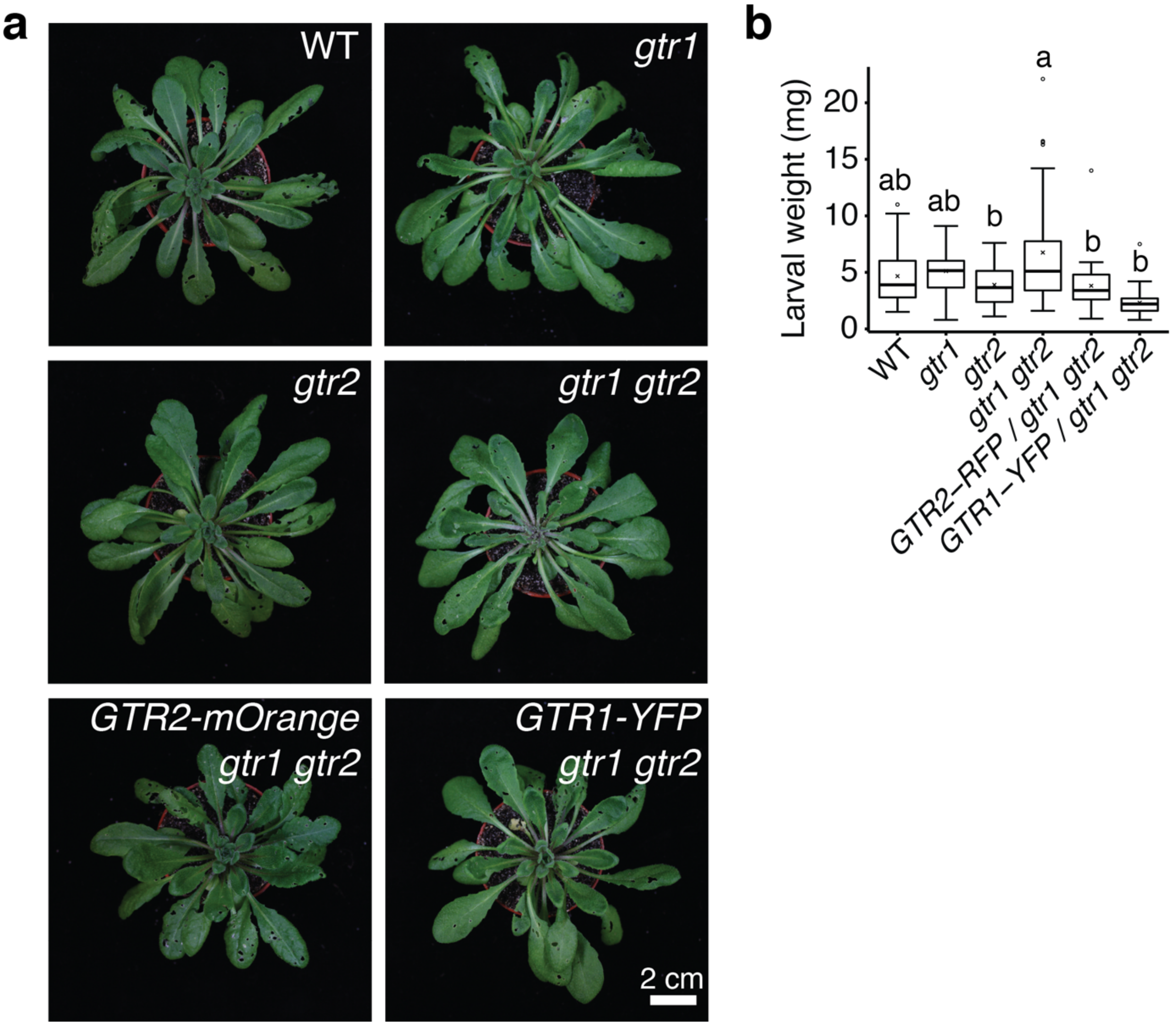


**Supplemental Fig. 2.** Complementation of the *S. littoralis* feeding preference phenotype of *gtr1gtr2*. **a**, Photographs of 5-week-old wild-type (WT), *gtr1*, *gtr2*, *gtr1 gtr2*, *pGTR2:GTR2-mOrange/gtr1 gtr2* and *pGTR1:GTR1-YFP/gtr1 gtr2* mutant plants imaged 7 days after infection with newly hatched larvae in a non-choice experimental set-up. Scale bar = 2 cm. **b**, Quantification of larval weight of *S. littoralis* following 7 days of feeding on 4-week-old wild-type (WT) and mutant plants. Boxplots show median (center line), mean (cross), first quartile (lower hinge), third quartile (upper hinge), whiskers (extending 1.5 times the inter-quartile range) and possible outliers (circles). Letters depict significant differences between genotypes (two-way ANOVA with post-hoc Tukey HSD test; n = 36, 22, 40, 34, 29, 33; p<0.001).

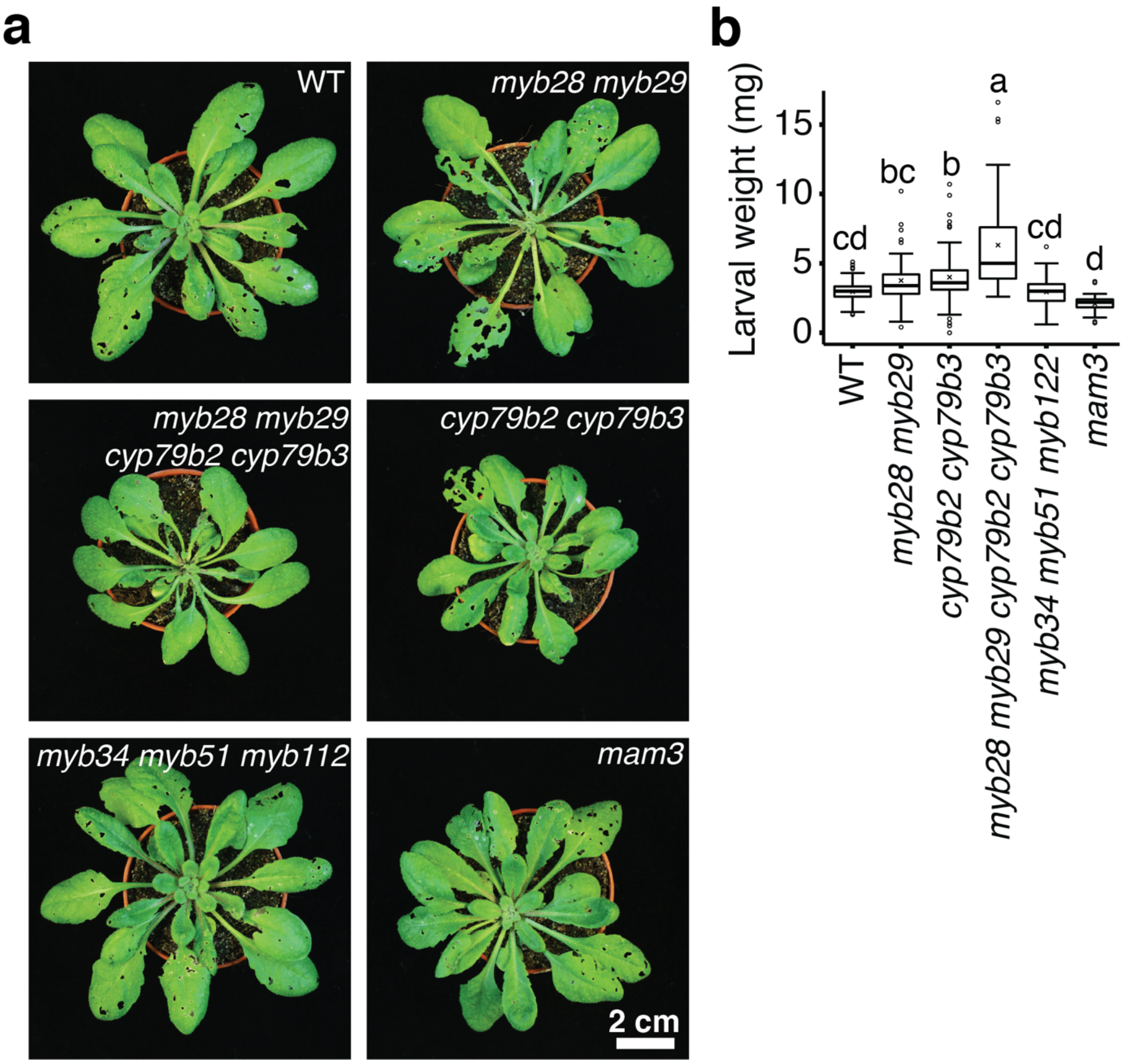


**Supplemental Fig. 3.** *S. littoralis* feeding preference on loss-of-function mutants for glucosinolate biosynthesis. **a**, Photographs of 4-week-old wild-type (WT), *myb28 myb29*, *cyp79b2 cyp79b3*, *myb28 myb29 cyp79b2 cyp79b3*, *myb34 myb51 myb112* and *mam3* mutant plants imaged 7 days after infection with newly hatched larvae in a non-choice experimental set-up. Scale bar = 2 cm. **b**, Quantification of larval weight of *S. littoralis* following 7 days of feeding on 4-week-old wild-type and mutant plants. Boxplots show median (center line), mean (cross), first quartile (lower hinge), third quartile (upper hinge), whiskers (extending 1.5 times the inter-quartile range) and possible outliers (circles). Letters depict significant differences between genotypes (two-way ANOVA with post-hoc Tukey HSD test; n = 61; 61; 60; 59; 59; 30; $p<0.001$). Pooled data of two independent replicates are shown.

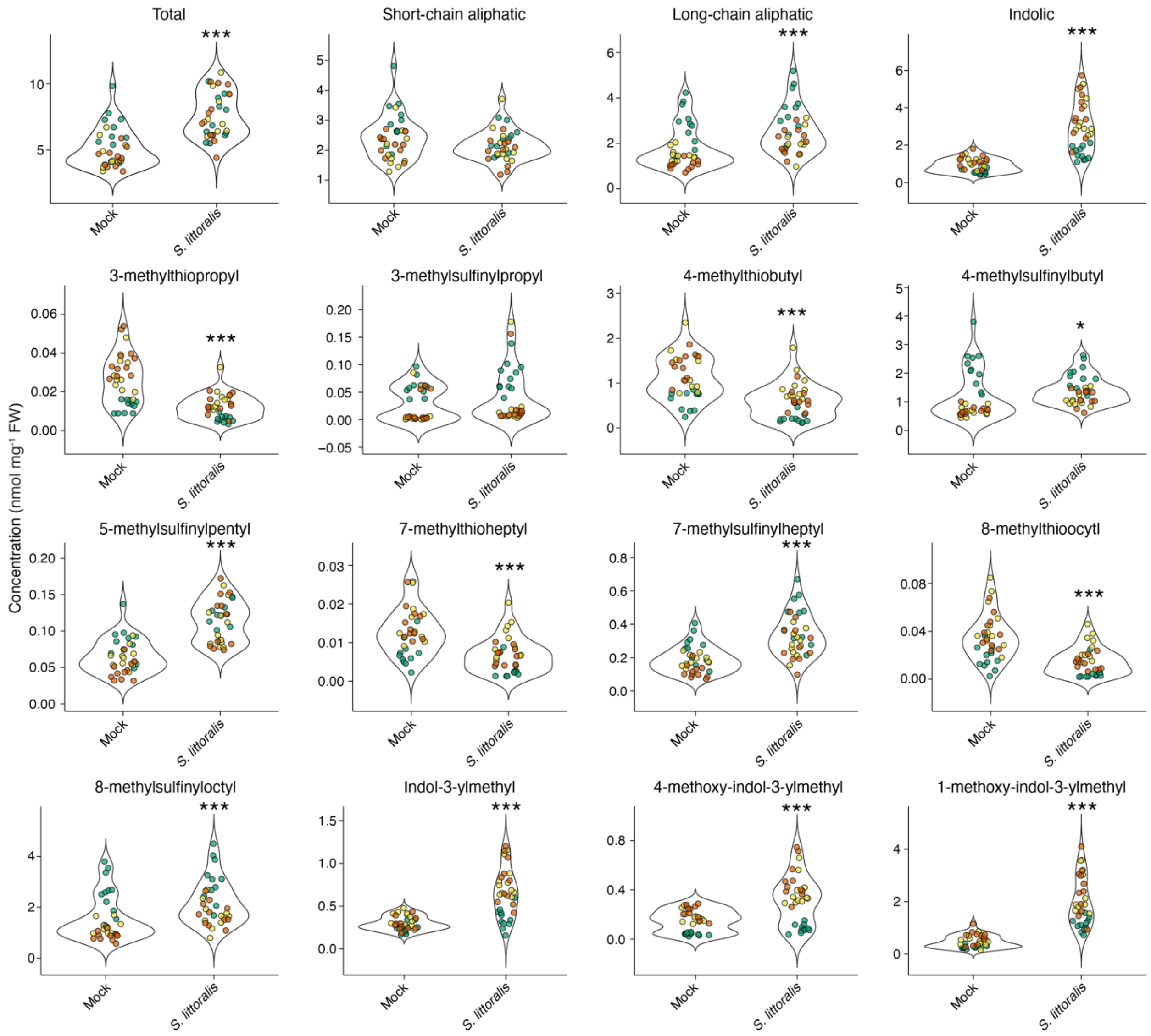


**Supplemental Fig. 4.** *S. littoralis*-induced perturbations of glucosinolate concentration in young leaves (L9+) of wild-type plants. Three newly hatched larvae per 5-week-old plant were allowed to feed for eight days. Young leaves were classified according to Fig. 1a. Asterisks indicate significant treatment effects (two-way ANOVA; n = 11; *p<0.05, **p<0.01, ***p<0.001). Pooled data from three independent experimental replicates are shown. Data points represent biological replicates and are color-coded according to experimental replicates.

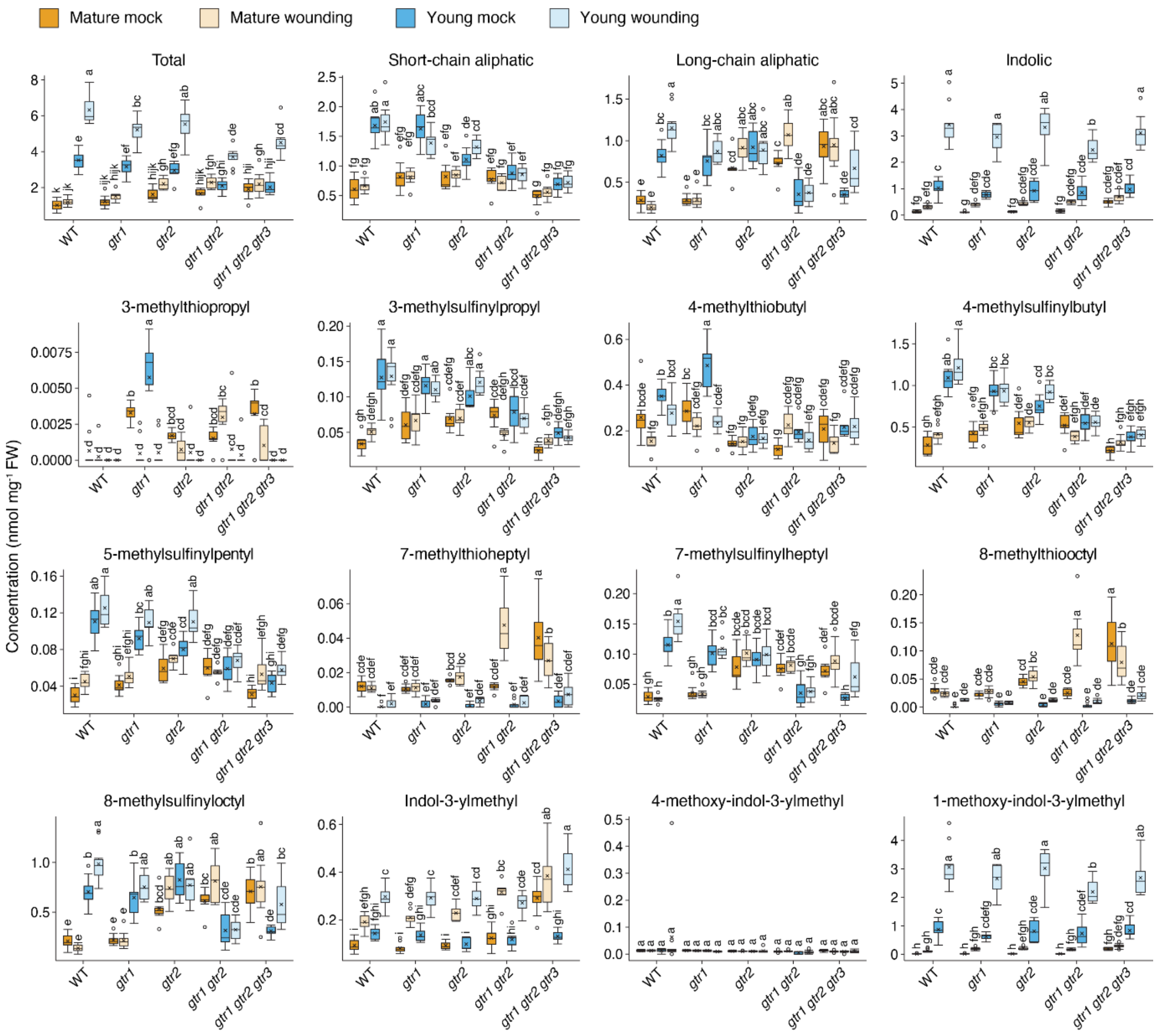


**Supplemental Fig. 5.** Tissue-specific glucosinolate analysis in mature (L6-8; orange) and young (L9+; blue) leaves of wild-type (WT), *gtr1*, *gtr2*, *gtr1 gtr2* and *gtr1 gtr2 gtr3* upon repetitive mechanical wounding of old leaves (L3-5). Samples were harvested 24 hours after treatment. Letters indicate significant differences of genotype × tissue interactions (two-way ANOVA; n = 10; p<0.001). Leaves were grouped according to Fig. 1a.

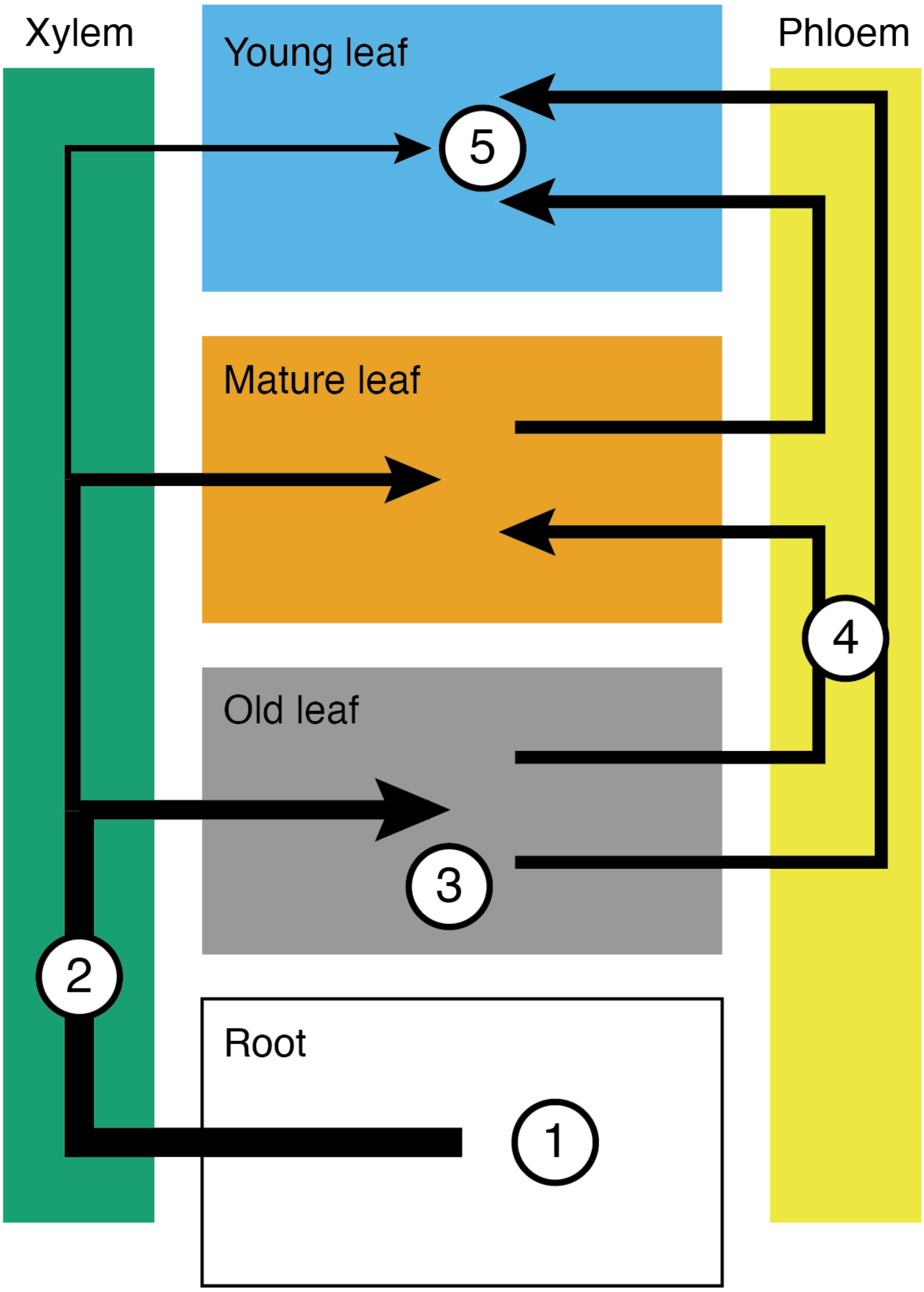


**Supplemental Fig. 6.** Model for glucosinolate transport upon herbivore attack. **(1)** GTR1-3 prevent root-synthesized glucosinolates from translocation to the shoot via the xylem under normal growth conditions providing a retention mechanism [19,38]. **(2)** Mechanical wounding of leaves by herbivore feeding attenuates GTR1/GTR2/GTR3-based glucosinolate retention in the root and leads to increased root-to-shoot translocation of mostly methylthioalkyl (3-methylthiopropyl, 4-methylthiobutyl, 7-methylthioheptyl and 8-methylthiooctyl) and indolic (indol-3-ylmethyl and 1-methoxy-indol-3-ylmethyl) glucosinolates via the xylem. Note that *GTR3* is exclusively expressed in roots [39]. **(3)** GTR1 and GTR2 load root-derived and old leaf-synthesized glucosinolates into the phloem of old leaves. High accumulation of glucosinolates in the extracellular space inhibits glucosinolate biosynthesis in old leaves. **(4)** Glucosinolates are translocated towards carbon sinks via the phloem [40–42]. **(5)** *S*-oxygenation of methylthioalkyl glucosinolates is induced either during transport in the phloem (i.e. inside the sieve element/companion cell-complex) or upon arrival in young leaves. Tissue-specific induction of *de novo* biosynthesis of indolic glucosinolates is independent of GTR1-3.

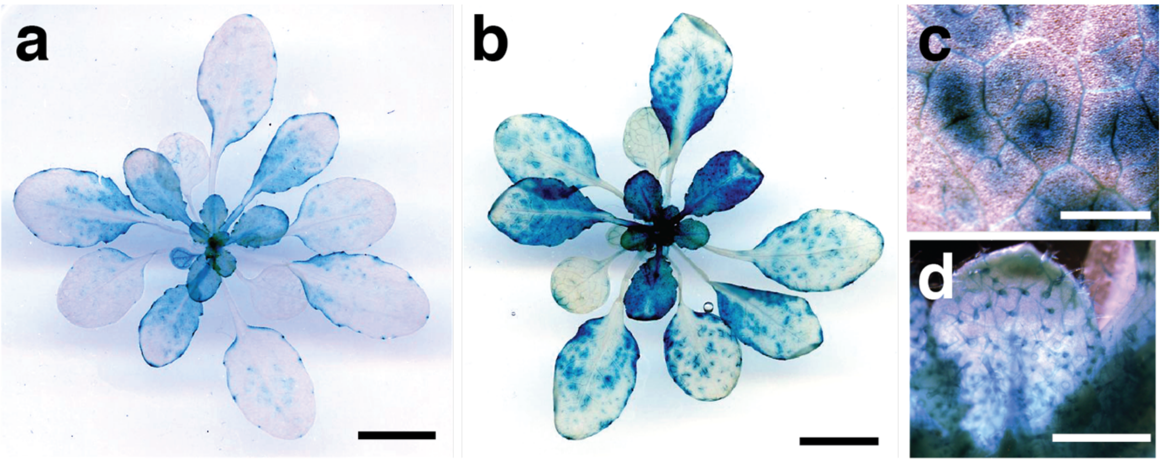


**Supplemental Fig. 7.** Gene expression patterns of *GTR1* and *GTR2* in 4-weeks-old plants. Scanning (**a-b**) and stereomicroscopic (**c-d**) images of GUS-stained transgenic plants that stably express *pGTR1::NLS-GUS-GFP* (**a**) and *pGTR2::NLS-GUS-GFP* (**b-d**). **c** and **d** represent higher magnification images of **b**. Scale bars = 1cm (a and b); 1 mm (c and d).

#### Supplemental Table 1. Oligonucleotides used for RT-qPCR analysis

| Gene alias | AGI code | Forward primer sequence | Reverse primer sequence |
|---|---|---|---|
| UBC21 | At5g25760 | CTGAGCCGGACAGTCCTCTTAACTG | CGGCGAGGCGTGTATACATTTGTG |
| MAM1 | At5g23010 | GCCGGTCAGTGTTACCCTTTCAATC | AGTTGCCGCATTTTTGGACACAG |
| MAM3 | At5g23020 | CGCTGATCTGAAGGCATTAGTGGTG | GCGGAAATCTGAGGGCTTGACATA |
| CYP83A1 | At4g13770 | TCTCGCCGCGGTTCTCCTTT | GCCCATCCAGCGAAGAAGCGT |
| CYP79B2 | At4g39950 | CCTGCGTGATCACTCCCTTT | TGTGTCTCGCTGGACATACG |
| SUR1 | At2g20610 | TGCGTAAAACTGGCCAGAGAGGA | ACCCAGAGCATCCCCTGCAA |
| UGT74C1 | At2g31790 | TGGACTTGTGGCTAAGTGGG | GCCTCCAATGTCGAGTTCCA |

### Supplemental Table 2. ANOVA tables for data shown in Figures

| Figure | Factor | *F* value | df | p-value |
|---|---|---|---|---|
| 1b | Genotype | 33.3082 | 1 | 8.513e-08 |
| | Tissue | 110.1216 | 2 | < 2.2e-16 |
| | Replicate | 1.5356 | 1 | 0.218117 |
| | Genotype × Tissue | 62.6424 | 2 | < 2.2e-16 |
| | Genotype × Replicate | 5.2599 | 1 | 0.023874 |
| | Tissue × Replicate | 2.8633 | 2 | 0.061678 |
| | Genotype × Tissue × Replicate | 4.8225 | 2 | 0.009972 |
| 1c – CYP79B2 | Genotype | 1.0588 | 1 | 0.317127 |
| | Tissue | 0.2847 | 2 | 0.755598 |
| | Genotype × Tissue | 6.6585 | 2 | 0.006846 |
| 1c – CYP83A1 | Genotype | 0.0853 | 1 | 0.77352 |
| | Tissue | 20.3946 | 2 | 2.365e-05 |
| | Genotype × Tissue | 4.7774 | 2 | 0.02166 |
| 1c – MAM1 | Genotype | 1.0361 | 1 | 0.3222 |
| | Tissue | 26.0209 | 2 | 4.889e-06 |
| | Genotype × Tissue | 1.6758 | 2 | 0.2151 |
| 1c – MAM3 | Genotype | 84.252 | 1 | 3.282e-08 |
| | Tissue | 143.805 | 2 | 8.530e-12 |
| | Genotype × Tissue | 46.832 | 2 | 7.349e-08 |
| 1c – SUR1 | Genotype | 18.2936 | 1 | 0.0004536 |
| | Tissue | 26.8889 | 2 | 3.922e-06 |
| | Genotype × Tissue | 7.8482 | 2 | 0.0035416 |
| 1c – UGT74C1 | Genotype | 5.1297 | 1 | 0.0361 |
| | Tissue | 3.8001 | 2 | 0.0420 |
| | Genotype × Tissue | 1.3912 | 2 | 0.2743 |
| 2c | Genotype | 346.4322 | 2 | < 2.2e-16 |
| | Replicate | 4.8403 | 2 | 0.008758 |
| | Genotype × Replicate | 14.1749 | 3 | 1.741e-08 |
| 3a – aliphatic | Treatment | 2.8467 | 1 | 0.09676 |
| | Replicate | 25.2868 | 2 | 1.084e-08 |
| | Treatment × Replicate | 0.6859 | 2 | 0.50752 |
| 3a – indolic | Treatment | 141.1673 | 1 | < 2.2e-16 |
| | Replicate | 26.0131 | 2 | 7.327e-09 |
| | Treatment × Replicate | 8.6047 | 2 | 0.0005181 |
| 3a – thio | Treatment | 49.4193 | 1 | 2.225e-09 |
| | Replicate | 35.4179 | 2 | 6.962e-11 |
| | Treatment × Replicate | 3.1494 | 2 | 0.05004 |
| 3a – sulfinyl | Treatment | 16.725 | 1 | 0.0001306 |
| | Replicate | 54.805 | 2 | 2.89e-14 |
| | Treatment × Replicate | 1.747 | 2 | 0.1830501 |
| 3b – aliphatic | Genotype | 20.3171 | 1 | 2.857e-05 |
| | Treatment | 4.6174 | 1 | 0.03544 |
| | Tissue | 116.1544 | 1 | 5.082e-16 |
| | Genotype × Treatment | 0.0838 | 1 | 0.77315 |
| | Genotype × Tissue | 207.5549 | 1 | < 2.2e-16 |
| | Treatment × Tissue | 0.4681 | 1 | 0.49633 |
| | Genotype × Treatment × Tissue | 5.2316 | 1 | 0.02550 |
| 3b – indolic | Genotype | 6.8209 | 1 | 0.0112121 |
| | Treatment | 173.9787 | 1 | < 2.2e-16 |
| | Tissue | 376.2752 | 1 | < 2.2e-16 |
| | Genotype × Treatment | 3.3115 | 1 | 0.0734753 |
| | Genotype × Tissue | 13.8237 | 1 | 0.0004247 |
| | Treatment × Tissue | 103.5383 | 1 | 5.294e-15 |
| | Genotype × Treatment × Tissue | 6.8880 | 1 | 0.0108394 |
| 3b – thio | Genotype | 37.952 | 1 | 5.374e-08 |
| | Treatment | 39.093 | 1 | 3.731e-08 |
| | Tissue | 122.985 | 1 | < 2.2e-16 |
| | Genotype × Treatment | 41.621 | 1 | 1.688e-08 |
| | Genotype × Tissue | 36.724 | 1 | 7.997e-08 |
| | Treatment × Tissue | 17.711 | 1 | 8.180e-05 |
| | Genotype × Treatment × Tissue | 48.081 | 1 | 2.428e-09 |
| 3b – sulfinyl | Genotype | 0.1888 | 1 | 0.6653791 |
| | Treatment | 9.6563 | 1 | 0.0028137 |
| | Tissue | 41.0030 | 1 | 2.046e-08 |
| | Genotype × Treatment | 0.0374 | 1 | 0.8473041 |
| | Genotype × Tissue | 260.5920 | 1 | < 2.2e-16 |
| | Treatment × Tissue | 3.1598 | 1 | 0.0802220 |
| | Genotype × Treatment × Tissue | 15.5010 | 1 | 0.0002061 |

### Supplemental Table 3. ANOVA tables for data shown in Supplemental Figures

| Figure | Factor | *F* value | df | p-value |
|---|---|---|---|---|
| 1 – total | Genotype | 33.3082 | 1 | 8.513e-08 |
| | Tissue | 110.1216 | 2 | < 2.2e-16 |
| | Replicate | 1.5356 | 1 | 0.218117 |
| | Genotype × Tissue | 62.6424 | 2 | < 2.2e-16 |
| | Genotype × Replicate | 5.2599 | 1 | 0.023874 |
| | Tissue × Replicate | 2.8633 | 2 | 0.061678 |
| | Genotype × Tissue × Replicate | 4.8225 | 2 | 0.009972 |
| 1 – indolic | Genotype | 5.0887 | 1 | 0.026217 |
| | Tissue | 147.6893 | 2 | < 2.2e-16 |
| | Replicate | 20.4952 | 1 | 1.624e-05 |
| | Genotype × Tissue | 0.1796 | 2 | 0.835836 |
| | Genotype × Replicate | 7.9320 | 1 | 0.005832 |
| | Tissue × Replicate | 20.1265 | 2 | 4.290e-08 |
| | Genotype × Tissue × Replicate | 4.0321 | 2 | 0.020637 |
| 1 – long-chain | Genotype | 110.4598 | 1 | < 2.2e-16 |
| | Tissue | 2.3099 | 2 | 0.104443 |
| | Replicate | 10.3306 | 1 | 0.001752 |
| | Genotype × Tissue | 111.3098 | 2 | < 2.2e-16 |
| | Genotype × Replicate | 5.0068 | 1 | 0.027423 |
| | Tissue × Replicate | 0.6498 | 2 | 0.524278 |
| | Genotype × Tissue × Replicate | 1.0103 | 2 | 0.367723 |
| 1 – short-chain | Genotype | 0.2342 | 1 | 0.629451 |
| | Tissue | 108.7709 | 2 | < 2.2e-16 |
| | Replicate | 3.9852 | 1 | 0.048567 |
| | Genotype × Tissue | 32.9098 | 2 | 9.368e-12 |
| | Genotype × Replicate | 0.2969 | 1 | 0.586999 |
| | Tissue × Replicate | 0.7982 | 2 | 0.452935 |
| | Genotype × Tissue × Replicate | 7.4634 | 2 | 0.000944 |
| 1 – 3mtp | Genotype | 25.7503 | 1 | 1.751e-06 |
| | Tissue | 22.3615 | 2 | 8.855e-09 |
| | Replicate | 134.2909 | 1 | < 2.2e-16 |
| | Genotype × Tissue | 1.1442 | 2 | 0.3225 |
| | Genotype × Replicate | 1.7593 | 1 | 0.1877 |
| | Tissue × Replicate | 36.1146 | 2 | 1.385e-12 |
| | Genotype × Tissue × Replicate | 0.9071 | 2 | 0.4069 |
| 1 – 3msp | Genotype | 4.9811 | 1 | 0.02781 |
| | Tissue | 92.5779 | 2 | < 2.2e-16 |
| | Replicate | 148.8662 | 1 | < 2.2e-16 |
| | Genotype × Tissue | 22.9322 | 2 | 5.964e-09 |
| | Genotype × Replicate | 0.6657 | 1 | 0.41645 |
| | Tissue × Replicate | 19.0964 | 2 | 9.028e-08 |
| | Genotype × Tissue × Replicate | 17.5191 | 2 | 2.882e-07 |
| 1 – 4mtb | Genotype | 0.0922 | 1 | 0.76203 |
| | Tissue | 46.9844 | 2 | 3.444e-15 |
| | Replicate | 56.6262 | 1 | 2.150e-11 |
| | Genotype × Tissue | 13.3068 | 2 | 7.328e-06 |
| | Genotype × Replicate | 0.7076 | 1 | 0.40220 |
| | Tissue × Replicate | 18.6235 | 2 | 1.275e-07 |
| | Genotype × Tissue × Replicate | 3.2476 | 2 | 0.04292 |
| 1 – 4msb | Genotype | 0.5063 | 1 | 0.4784 |
| | Tissue | 136.2459 | 2 | < 2.2e-16 |
| | Replicate | 106.6884 | 1 | < 2.2e-16 |
| | Genotype × Tissue | 44.7268 | 2 | 1.131e-14 |
| | Genotype × Replicate | 0.0971 | 1 | 0.7560 |
| | Tissue × Replicate | 11.5076 | 2 | 3.115e-05 |
| | Genotype × Tissue × Replicate | 26.1300 | 2 | 6.882e-10 |
| 1 – 5msp | Genotype | 6.9692 | 1 | 0.009595 |
| | Tissue | 83.3707 | 2 | < 2.2e-16 |
| | Replicate | 212.0631 | 1 | < 2.2e-16 |
| | Genotype × Tissue | 49.7094 | 2 | 8.502e-16 |
| | Genotype × Replicate | 0.5975 | 1 | 0.441314 |
| | Tissue × Replicate | 22.5003 | 2 | 8.041e-09 |
| | Genotype × Tissue × Replicate | 34.1299 | 2 | 4.486e-12 |
| 1 – 7mth | Genotype | 110.0477 | 1 | < 2.2e-16 |
| | Tissue | 38.5960 | 2 | 3.306e-13 |
| | Replicate | 51.7426 | 1 | 1.090e-10 |
| | Genotype × Tissue | 45.9344 | 2 | 5.966e-15 |
| | Genotype × Replicate | 19.5089 | 1 | 2.499e-05 |
| | Tissue × Replicate | 1.5067 | 2 | 0.2265348 |
| | Genotype × Tissue × Replicate | 8.9770 | 2 | 0.0002564 |
| 1 – 7msh | Genotype | 56.5616 | 1 | 2.196e-11 |
| | Tissue | 8.9357 | 2 | 0.0002655 |
| | Replicate | 39.6398 | 1 | 7.782e-09 |
| | Genotype × Tissue | 133.1091 | 2 | < 2.2e-16 |
| | Genotype × Replicate | 0.0005 | 1 | 0.9813406 |
| | Tissue × Replicate | 2.3814 | 2 | 0.0975383 |
| | Genotype × Tissue × Replicate | 19.1839 | 2 | 8.472e-08 |
| 1 – 8mto | Genotype | 75.729 | 1 | 5.988e-14 |
| | Tissue | 15.354 | 2 | 1.481e-06 |
| | Replicate | 107.415 | 1 | < 2.2e-16 |
| | Genotype × Tissue | 31.269 | 2 | 2.564e-11 |
| | Genotype × Replicate | 32.759 | 1 | 1.053e-07 |
| | Tissue × Replicate | 1.073 | 2 | 0.3458 |
| | Genotype × Tissue × Replicate | 14.541 | 2 | 2.779e-06 |

| Figure | Factor | *F* value | df | p-value |
|---|---|---|---|---|
| 1 – 8mso | Genotype | 104.2332 | 1 | < 2.2e-16 |
| | Tissue | 6.0649 | 2 | 0.003245 |
| | Replicate | 7.0970 | 1 | 0.008976 |
| | Genotype × Tissue | 110.3851 | 2 | < 2.2e-16 |
| | Genotype × Replicate | 2.7319 | 1 | 0.101439 |
| | Tissue × Replicate | 0.5973 | 2 | 0.552209 |
| | Genotype × Tissue × Replicate | 0.7834 | 2 | 0.459559 |
| 1 – I3M | Genotype | 31.1046 | 1 | 2.015e-07 |
| | Tissue | 5.7502 | 2 | 0.004303 |
| | Replicate | 0.0945 | 1 | 0.759161 |
| | Genotype × Tissue | 6.3732 | 2 | 0.002466 |
| | Genotype × Replicate | 6.3087 | 1 | 0.013583 |
| | Tissue × Replicate | 0.5773 | 2 | 0.563213 |
| | Genotype × Tissue × Replicate | 0.6772 | 2 | 0.510304 |
| 1 – 4MOI3M | Genotype | 30.2820 | 1 | 2.791e-07 |
| | Tissue | 14.6448 | 2 | 2.564e-06 |
| | Replicate | 40.3099 | 1 | 6.083e-09 |
| | Genotype × Tissue | 0.9923 | 2 | 0.37429 |
| | Genotype × Replicate | 6.4596 | 1 | 0.01254 |
| | Tissue × Replicate | 1.3920 | 2 | 0.25326 |
| | Genotype × Tissue × Replicate | 4.6467 | 2 | 0.01171 |
| 1 – 1MOI3M | Genotype | 1.1002 | 1 | 0.29670 |
| | Tissue | 156.2959 | 2 | < 2.2e-16 |
| | Replicate | 22.9502 | 1 | 5.658e-06 |
| | Genotype × Tissue | 0.1940 | 2 | 0.82398 |
| | Genotype × Replicate | 5.2183 | 1 | 0.02442 |
| | Tissue × Replicate | 24.8310 | 2 | 1.636e-09 |
| | Genotype × Tissue × Replicate | 3.5525 | 2 | 0.03225 |
| 2b | Genotype | 8.9094 | 5 | 1.305e-07 |
| 3b | Genotype | 34.406 | 6 | <2.2e-16 |
| | Replicate | 83.619 | 1 | <2.2e-16 |
| | Genotype × Replicate | 14.797 | 5 | 2.756e-13 |
| 4 – total | Treatment | 44.2263 | | 9.85e-09 |
| | Replicate | 1.9508 | | 0.1511 |
| | Treatment × Replicate | 3.7697 | | 0.0287 |
| 4 – indolic | Treatment | 141.1673 | | <2.20e-16 |
| | Replicate | 26.0131 | | 7.33e-09 |
| | Treatment × Replicate | 8.6047 | | 0.0005181 |
| 4 – long-chain | Treatment | 20.8117 | | 2.56e-05 |
| | Replicate | 40.1347 | | 8.62e-12 |
| | Treatment × Replicate | 0.9095 | | 0.4082 |
| 4 – short-chain | Treatment | 3.4734 | | 0.067257 |
| | Replicate | 7.065 | | 0.001757 |
| | Treatment × Replicate | 2.3757 | | 0.101636 |
| 4 – 3mtp | Treatment | 77.829 | 1 | 1.97e-12 |
| | Replicate | 38.146 | 2 | 2.04e-11 |
| | Treatment × Replicate | 9.148 | 2 | 0.0003407 |
| 4 – 3msp | Treatment | 1.9647 | 1 | 0.1662 |
| | Replicate | 14.4194 | 2 | 7.70e-06 |
| | Treatment × Replicate | 0.1719 | 2 | 0.8425 |
| 4 – 4mtb | Treatment | 47.5296 | 1 | 3.79e-09 |
| | Replicate | 34.9539 | 2 | 8.62e-11 |
| | Treatment × Replicate | 3.1092 | 2 | 0.0519 |
| 4 – 4msb | Treatment | 4.7049 | 1 | 0.034049 |
| | Replicate | 68.3237 | 2 | 3.42e-16 |
| | Treatment × Replicate | 6.5874 | 2 | 0.002592 |
| 4 – 5msp | Treatment | 69.2901 | 1 | 1.38e-11 |
| | Replicate | 5.0083 | 2 | 0.009739 |
| | Treatment × Replicate | 2.8509 | 2 | 0.065647 |
| 4 – 7mth | Treatment | 36.0334 | 1 | 1.22e-07 |
| | Replicate | 23.4636 | 2 | 2.96e-08 |
| | Treatment × Replicate | 1.4497 | 2 | 0.2427 |
| 4 – 7msh | Treatment | 44.073 | 1 | 1.03e-08 |
| | Replicate | 14.232 | 2 | 8.74e-06 |
| | Treatment × Replicate | 1.136 | 2 | 0.3279 |
| 4 – 8mto | Treatment | 43.312 | 1 | 1.29e-08 |
| | Replicate | 22.4641 | 2 | 5.22e-08 |
| | Treatment × Replicate | 1.7018 | 2 | 0.191 |
| 4 – 8mso | Treatment | 18.5976 | 1 | 6.11e-05 |
| | Replicate | 45.2546 | 2 | 1.04e-12 |
| | Treatment × Replicate | 1.0038 | 2 | 0.3726 |
| 4 – I3M | Treatment | 89.251 | 1 | 1.76e-13 |
| | Replicate | 21.415 | 2 | 9.57e-08 |
| | Treatment × Replicate | 10.505 | 2 | 0.0001225 |
| 4 – 4MOI3M | Treatment | 64.8947 | 1 | 3.94e-11 |
| | Replicate | 68.53 | 2 | 3.21e-16 |
| | Treatment × Replicate | 8.8659 | 2 | 0.0004232 |
| 4 – 1MOI3M | Treatment | 134.9074 | 1 | <2.20e-16 |
| | Replicate | 16.6866 | 2 | 1.73e-06 |
| | Treatment × Replicate | 6.1222 | 2 | 0.003805 |
| 5 – total | Genotype | 8.5222 | 4 | 2.89e-06 |
| | Tissue | 860.2999 | 1 | <2.20e-16 |
| | Treatment | 299.9755 | 1 | <2.20e-16 |
| | Genotype × Tissue | 47.9588 | 4 | <2.20e-16 |
| | Genotype × Treatment | 1.2041 | 4 | 0.31113 |
| | Tissue × Treatment | 156.5235 | 1 | <2.20e-16 |
| | Genotype × Tissue × Treatment | 2.9453 | 4 | 0.02202 |

| Figure | Factor | *F* value | df | p-value |
|---|---|---|---|---|
| 5 – indolic | Genotype | 4.9163 | 4 | 0.000911 |
| | Tissue | 991.6936 | 1 | <2.20e-16 |
| | Treatment | 528.3926 | 1 | <2.20e-16 |
| | Genotype × Tissue | 5.2871 | 4 | 0.0004997 |
| | Genotype × Treatment | 1.4112 | 4 | 0.2325435 |
| | Tissue × Treatment | 322.3318 | 1 | <2.20e-16 |
| | Genotype × Tissue × Treatment | 2.134 | 4 | 0.0789427 |
| 5 – long-chain | Genotype | 12.4071 | 4 | 7.95e-09 |
| | Tissue | 14.4922 | 1 | 0.0001993 |
| | Treatment | 19.7901 | 1 | 1.60e-05 |
| | Genotype × Tissue | 82.2889 | 4 | <2.20e-16 |
| | Genotype × Treatment | 0.6065 | 4 | 0.6585145 |
| | Tissue × Treatment | 2.01 | 1 | 0.1581831 |
| | Genotype × Tissue × Treatment | 6.6831 | 4 | 5.29e-05 |
| 5 – short-chain | Genotype | 60.7359 | 4 | <2.20e-16 |
| | Tissue | 321.3976 | 1 | <2.20e-16 |
| | Treatment | 0.136 | 1 | 0.71279 |
| | Genotype × Tissue | 39.2784 | 4 | <2.20e-16 |
| | Genotype × Treatment | 2.1491 | 4 | 0.07712 |
| | Tissue × Treatment | 0.1881 | 1 | 0.66505 |
| | Genotype × Tissue × Treatment | 1.3887 | 4 | 0.24015 |
| 5 – 3mtp | Genotype | 17.871 | 4 | 3.58e-12 |
| | Tissue | 15.5267 | 1 | 0.0001207 |
| | Treatment | 41.0613 | 1 | 1.54e-09 |
| | Genotype × Tissue | 10.5371 | 4 | 1.30e-07 |
| | Genotype × Treatment | 18.0708 | 4 | 2.73e-12 |
| | Tissue × Treatment | 0.324 | 1 | 0.5699966 |
| | Genotype × Tissue × Treatment | 5.2837 | 4 | 0.0005025 |
| 5 – 3msp | Genotype | 45.5923 | 4 | <2.20e-16 |
| | Tissue | 202.5642 | 1 | <2.20e-16 |
| | Treatment | 0.2129 | 1 | 0.6451 |
| | Genotype × Tissue | 21.1473 | 4 | 4.79e-14 |
| | Genotype × Treatment | 2.7184 | 4 | 0.0316 |
| | Tissue × Treatment | 0.4893 | 1 | 0.4853 |
| | Genotype × Tissue × Treatment | 1.9698 | 4 | 0.1015 |
| 5 – 4mtb | Genotype | 36.5303 | 4 | <2.20e-16 |
| | Tissue | 44.2808 | 1 | 4.18e-10 |
| | Treatment | 36.2645 | 1 | 1.12e-08 |
| | Genotype × Tissue | 5.4734 | 4 | 0.0003697 |
| | Genotype × Treatment | 14.3052 | 4 | 5.07e-10 |
| | Tissue × Treatment | 6.9908 | 1 | 0.0089982 |
| | Genotype × Tissue × Treatment | 7.6111 | 4 | 1.21e-05 |
| 5 – 4msb | Genotype | 67.7619 | 4 | <2.20e-16 |
| | Tissue | 371.2701 | 1 | <2.20e-16 |
| | Treatment | 6.8798 | 1 | 0.00955 |
| | Genotype × Tissue | 46.6354 | 4 | <2.20e-16 |
| | Genotype × Treatment | 2.4815 | 4 | 0.04593 |
| | Tissue × Treatment | 0.3119 | 1 | 0.57726 |
| | Genotype × Tissue × Treatment | 1.5875 | 4 | 0.18008 |
| 5 – 5msp | Genotype | 38.1903 | 4 | <2.20e-16 |
| | Tissue | 331.6435 | 1 | <2.20e-16 |
| | Treatment | 45.2482 | 1 | 2.84e-10 |
| | Genotype × Tissue | 48.3326 | 4 | <2.20e-16 |
| | Genotype × Treatment | 2.0914 | 4 | 0.0843 |
| | Tissue × Treatment | 1.854 | 1 | 0.1752 |
| | Genotype × Tissue × Treatment | 1.3755 | 4 | 0.2447 |
| 5 – 7mth | Genotype | 24.7863 | 4 | 5.07e-16 |
| | Tissue | 264.0617 | 1 | <2.20e-16 |
| | Treatment | 7.3902 | 1 | 0.007273 |
| | Genotype × Tissue | 16.323 | 4 | 2.97e-11 |
| | Genotype × Treatment | 12.3842 | 4 | 8.22e-09 |
| | Tissue × Treatment | 0.3913 | 1 | 0.532491 |
| | Genotype × Tissue × Treatment | 15.119 | 4 | 1.60e-10 |
| 5 – 7msh | Genotype | 15.1481 | 4 | 1.53e-10 |
| | Tissue | 63.2216 | 1 | 2.99e-13 |
| | Treatment | 19.3114 | 1 | 2.00e-05 |
| | Genotype × Tissue | 93.9557 | 4 | <2.20e-16 |
| | Genotype × Treatment | 1.9792 | 4 | 0.1001 |
| | Tissue × Treatment | 3.9312 | 1 | 0.04909 |
| | Genotype × Tissue × Treatment | 2.4817 | 4 | 0.04592 |
| 5 – 8mto | Genotype | 31.325 | 4 | <2.20e-16 |
| | Tissue | 253.6411 | 1 | <2.20e-16 |
| | Treatment | 11.8919 | 1 | 0.0007186 |
| | Genotype × Tissue | 20.636 | 4 | 9.25e-14 |
| | Genotype × Treatment | 14.9859 | 4 | 1.93e-10 |
| | Tissue × Treatment | 0.4961 | 1 | 0.4822263 |
| | Genotype × Tissue × Treatment | 16.1581 | 4 | 3.73e-11 |
| 5 – 8mso | Genotype | 12.2 | 4 | 1.08e-08 |
| | Tissue | 35.3563 | 1 | 1.64e-08 |
| | Treatment | 16.549 | 1 | 7.39e-05 |
| | Genotype × Tissue | 73.9592 | 4 | <2.20e-16 |
| | Genotype × Treatment | 0.5443 | 4 | 0.7034306 |
| | Tissue × Treatment | 2.24 | 1 | 0.1364261 |
| | Genotype × Tissue × Treatment | 5.5166 | 4 | 0.0003448 |

| **Figure** | **Factor** | ***F* value** | **df** | **p-value** |
|---|---|---|---|---|
| 5 – I3M | Genotype | 44.0054 | 4 | <2.20e-16 |
| | Tissue | 6.2914 | 1 | 0.0131164 |
| | Treatment | 429.4863 | 1 | <2.20e-16 |
| | Genotype × Tissue | 14.3232 | 4 | 4.94e-10 |
| | Genotype × Treatment | 2.1995 | 4 | 0.0713322 |
| | Tissue × Treatment | 16.6232 | 1 | 7.13e-05 |
| | Genotype × Tissue × Treatment | 5.9804 | 4 | 0.0001632 |
| 5 – 4MOI3M | Genotype | 1.446 | 4 | 0.2212 |
| | Tissue | 0.6398 | 1 | 0.4249 |
| | Treatment | 0.9806 | 1 | 0.3235 |
| | Genotype × Tissue | 1.1351 | 4 | 0.3419 |
| | Genotype × Treatment | 1.0128 | 4 | 0.4025 |
| | Tissue × Treatment | 1.2053 | 1 | 0.2739 |
| | Genotype × Tissue × Treatment | 0.9758 | 4 | 0.4224 |
| 5 – 1MOI3M | Genotype | 3.4029 | 4 | 0.010566 |
| | Tissue | 1024.9995 | 1 | <2.20e-16 |
| | Treatment | 421.3365 | 1 | <2.20e-16 |
| | Genotype × Tissue | 3.5748 | 4 | 0.008007 |
| | Genotype × Treatment | 1.7306 | 4 | 0.145693 |
| | Tissue × Treatment | 316.6292 | 1 | <2.20e-16 |
| | Genotype × Tissue × Treatment | 1.6772 | 4 | 0.157763 |
